# Stable Network Motifs with Divergent Engagement Across Memory Outcomes

**DOI:** 10.64898/2026.09.19.752867

**Authors:** Aditya Kumar, Kevin Tyner, Ross Moseley, Matthew Svalina, Aaron S. Geller, Ueli Rutishauser, Daniel R. Kramer, John A. Thompson

## Abstract

Visual recognition memory depends on coordinated interactions across distributed medial temporal, prefrontal, and limbic networks, yet the sub-second large-scale network dynamics distinguishing successful from unsuccessful recognition remain poorly understood. Here we used intracranial EEG recordings from seven patients (4 female, 3 male) with medically refractory epilepsy performing a new–old visual recognition memory task to characterize time-resolved directed functional connectivity and dynamic network organization. Directed effective connectivity was estimated using bivariate Granger causality in a sliding-window framework, and dynamic community structure was identified using the Louvain modularity algorithm. Successful recognition (Hits) was associated with a qualitatively distinct network configuration: low modularity (Q = 0.05 vs. 0.09–0.15 in other conditions), a cross-hemispheric community structure integrating left medial temporal lobe (MTL) with right prefrontal cortex, and sustained network integration throughout the post-stimulus period. Right-hemisphere nodes within this cross-hemispheric community showed stronger within-community directed connectivity than left-hemisphere nodes (mean asymmetry index = 0.33, p < 0.001), consistent with right prefrontal dominance in driving network integration during successful retrieval. In contrast, Correct Rejections were characterized by early transient network segregation (onset 32 ms) dominated by right-hemisphere activity. Hub regions were non-overlapping across conditions—left-lateralized during Hits, right-lateralized during Correct Rejections. These findings imply that successful visual recognition is distinguished not by stronger connectivity per se, but by a reorganization of large-scale network topology that transiently integrates left MTL with right prefrontal cortex into a unified functional community.

**Significance Statement:** How does the brain recognize something it has seen before? This study used direct intracranial brain recordings in epilepsy patients to examine how networks of brain regions communicate during successful and unsuccessful recognition memory. We found that successful recognition is not simply a matter of greater connectivity strength. It requires a qualitatively different pattern of large-scale network organization in which left-hemisphere memory regions and right-hemisphere evaluation regions form a unified, cross-hemispheric network. Unsuccessful rejection of novel images, by contrast, was characterized by early, segregated right-hemisphere activity. These findings therefore present a potential, specific, circuit-level mechanism supporting recognition memory, with implications for understanding memory dysfunction in neurological disorders such as epilepsy and Alzheimer’s disease.

## Introduction

Visual recognition, the ability to identify whether a stimulus was previously encountered, is a fundamental component of declarative memory (Milner 1972, Squire, Wixted et al. 2007). Behavioral work has demonstrated that visual recognition is remarkably accurate, rapid, and durable over long delays of days to years (Standing 1973, Brady, Konkle et al. 2008, Besson, Ceccaldi et al. 2012, Larzabal, Tramoni et al. 2018). Neurological studies have consistently implicated medial temporal lobe (MTL) structures (Brown and Aggleton 2001, Eichenbaum, Yonelinas et al. 2007, Staresina, Reber et al. 2019) as critical for successful recognition. Lesion, stimulation, and single-unit studies converge on the importance of these structures for visual recognition memory (Milner 1972, Quiroga, Reddy et al. 2005, Squire, Wixted et al. 2007, Selimbeyoglu and Parvizi 2010). However, recognition memory is not supported by the MTL in isolation. Rather, it emerges from interactions among distributed cortical and subcortical regions spanning temporal, frontal, parietal, and limbic networks (Rugg and Vilberg 2013, Wynn and Nyhus 2022). How the communication between these networks is organized at the sub-second timescale remains poorly understood, limiting our understanding of the circuit-level mechanisms that support or fail recognition.

Large-scale brain networks exhibit dynamic, non-random organization that reconfigures rapidly in response to changing cognitive demands (Bassett, Wymbs et al. 2011, Bullmore and Sporns 2012). These reconfigurations have been increasingly linked to successful memory performance (Cohen and D’Esposito 2016). Although overt behavioral responses typically occur several hundred milliseconds after stimulus onset (Besson, Ceccaldi et al. 2012), neural signatures differentiating previously-seen from novel stimuli emerge as early as ∼200 ms (Barbeau, Taylor et al. 2008, Barragan-Jason, Cauchoix et al. 2015). These effects are followed by sustained activity across multiple regions extending well into post-stimulus intervals associated with decision formation and memory evaluation (Rugg and Curran 2007, López-Madrona, Trébuchon et al. 2024).

Despite this progress, a gap remains in understanding the sub-second network dynamics of recognition memory. Many studies of large-scale connectivity rely on fMRI, which offers high spatial but limited temporal resolution, or scalp EEG/MEG, which offers the reverse (Hutchison, Womelsdorf et al. 2013). Intracranial EEG (iEEG) overcomes these limitations by providing direct recordings of neural activity with millisecond resolution and precise localization (Lachaux, Axmacher et al. 2012). Recent iEEG studies have characterized rapid reorganization of effective connectivity and network topology within the first few hundred milliseconds after stimulus presentation (Kopal, Hlinka et al. 2023).

Several important questions remain unresolved. First, it is unclear how directed functional interactions differ between successful recognition of previously encountered stimuli and correct rejection of novel stimuli. Second, less is known about how network segregation and modular organization differ as a function of recognition outcome (Das and Menon 2024). Finally, it is unclear whether successful recognition is associated with selectively strengthened directed interactions along specific anatomical pathways, and whether these pathways are shared with those supporting correct rejection.

We addressed these questions by examining time-resolved directed functional connectivity during a new–old visual recognition memory task using iEEG recordings from patients with medically refractory epilepsy. Directed interactions were quantified using bivariate Granger causality, a measure of asymmetric predictability between neural signals (Granger 1969, Ding, Chen et al. 2006, Seth, Barrett et al. 2015). By combining sliding-window Granger causality with graph-theoretic and information-theoretic analyses, we characterized the dynamic evolution of large-scale brain networks spanning medial temporal, anterior cingulate, and orbitofrontal regions in both hemispheres.

Here, we report that successful recognition of a previously identified stimulus (Hits) is associated with greater magnitude of directed functional connectivity relative to correct rejection (CR) of novel stimuli (correctly identifying the stimulus as unseen), particularly during post-stimulus intervals. We further report that hits exhibit distinct network topology, including community structure and hub organization, compared to CR. By linking time-resolved directed connectivity, network organization, and behavioral outcome, this work aims to elucidate the dynamic large-scale mechanisms that support successful visual recognition memory.

## Methods

### Participants

Participants were adult patients with medically refractory epilepsy undergoing clinical monitoring. Inclusion criteria were: (1) intractable epilepsy undergoing invasive monitoring; (2) age ≥13; (3) Full Scale Intelligence Quotient >70; and (4) ability to comprehend and perform simple behavioral tasks by pressing buttons on a laptop computer in response to questions. Exclusion criteria were: determination by clinicians and investigators that a patient was unable to complete the behavioral tasks required for the protocol due to cognitive limits, psychological limits, or pain.

Seven patients (ages 23–49; 3 male, 4 female) with medically refractory epilepsy undergoing clinical monitoring for seizure localization were enrolled. All participants were screened for cognitive deficits by a clinical neuropsychologist and were excluded if they had cognitive or memory impairment. Participants were implanted with intracranial electrodes for clinical purposes with locations chosen by the clinical team, and were recruited to perform a cognitive memory task during their hospital stay. All procedures were approved by the COMIRB (CEDAR IRB). Written informed consent was obtained from all participants prior to data collection.

### New-old memory delay experimental design and task

Participants performed a new–old recognition memory task consisting of a learning portion followed by a recognition portion within the same recording session (Kopal, Hlinka et al. 2023). Stimuli were presented on a monitor positioned in front of the patient, constrained by the hospital room setup. The task was implemented using Psychtoolbox 3.0 on MATLAB R2022a (Brainard 1997, Pelli 1997, Kleiner, Brainard et al. 2007, The MathWorks 2022). During the learning portion (Learn), participants were sequentially presented with 100 images, each presented only once, drawn from several stimulus categories (e.g., objects, animals, landscapes). Participants were instructed to attend to each image in preparation for a subsequent memory test (e.g., answer whether the presented image contained an animal).

After a delay period of experimenter-determined variable duration across participants (approximately 5 minutes maximum), subjects completed the recognition portion of the task. In this portion 100 images were presented, consisting of 50 previously seen (“old”) images and 50 novel (“new”) images. Trial order was pseudo-randomized.

Each learn trial followed a fixed structure: an intertrial interval (ITI, 1s) displaying a gray screen, followed by stimulus presentation (1s), a response screen (self-paced), and a subsequent ITI preceding the next trial. During the response period, participants indicated whether they believed the image presented did or did not represent an “animal.” Feedback regarding task performance was not provided (Figure 1).

**Figure 1.**
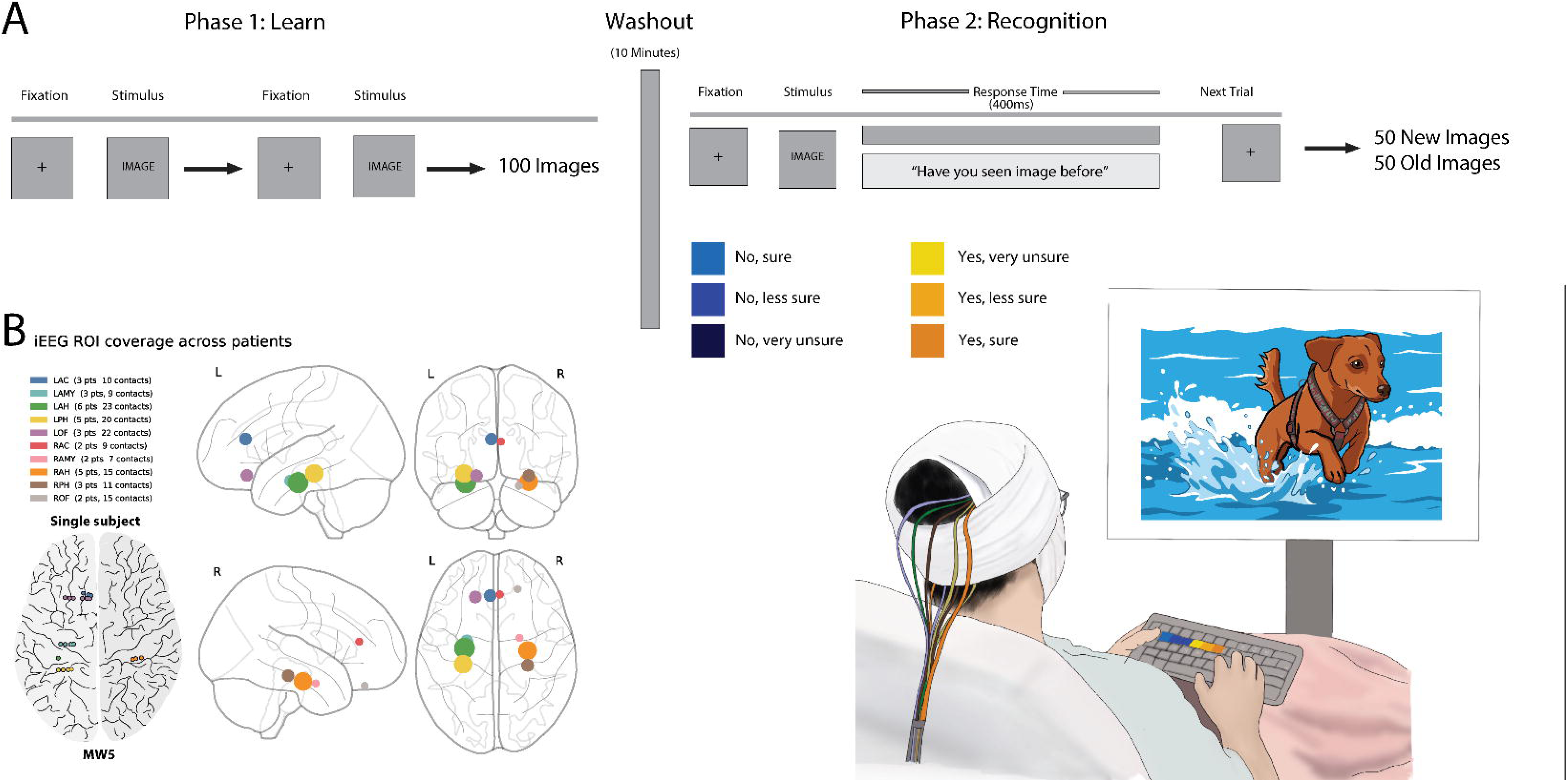
Task Depiction and Electrode Coverage. (A) An artist’s depiction of the task and intra-task timeline (B) ROI determined electrode coverage. Marker color indicates ROI and marker size represents number of total contacts within respective ROI.

Similarly, each recognition trial followed a fixed structure: an intertrial interval (ITI, 1s) displaying a gray screen, followed by stimulus presentation (1s), a response screen (self-paced), and a subsequent ITI preceding the next trial. During the response period, participants indicated whether they believed the image had been previously seen or was new and provided a confidence rating for their judgment (1–6 confidence scale using a computer keyboard). Feedback regarding task performance was not provided (Figure 1).

Behavioral responses were categorized based on recognition accuracy. Hits were defined as trials in which previously presented images were correctly identified as old, and Correct Rejections (CRs) were defined as trials in which novel images were correctly identified as new (Kopal, Hlinka et al. 2023). Although neural and behavioral data from Misses (when patients saw the image during encoding but failed to recognize later) and False Alarms (when patients did not see the image during encoding but believed they had later) were available, these trial types were not a central focus of the current study and were reserved for supplementary analyses.

Recognition memory performance was quantified using signal detection theory (Macmillan and Creelman 2005), which separates sensitivity (the ability to discriminate old from new items) from response bias (the tendency to favor “old” or “new” responses). Sensitivity (d’) was computed as the difference between the z-transformed hit rate and z-transformed false alarm rate. Response bias (criterion, c) was computed as the negative average of the z-transformed hit rate and false alarm rate (c = −0.5 x [z(H) + z(FA)]). This measure reflects the participant’s decision threshold, with positive values indicating a conservative bias (tendency to respond “new”) and negative values indicating a liberal bias (tendency to respond “old”). This formulation allows sensitivity and bias to be estimated independently, which is critical for interpreting recognition performance.

### IEEG data acquisition and channel selection

#### Recording Equipment and Data Acquisition

Intracranial EEG signals were recorded using a Neuralynx Atlas clinical recording system (Neuralynx, Inc., Bozeman, MT). Electrodes were stereotactically implanted depth electrodes (PMT Corporation, Chanhassen, MN, or DIXI Medical, Besançon, France; both standard clinical stereo-EEG macroelectrodes), with number, trajectory, and contact spacing determined entirely by the clinical team based on the hypothesized seizure onset zone. Continuous data were sampled at 4 kHz and subsequently downsampled to 500 Hz for all offline analyses described below.

#### Electrode Localization and Channel Selection

To facilitate group-level analyses while accommodating heterogeneous electrode coverage, analyses were restricted to a predefined set of anatomically defined regions of interest (ROIs). Study participants contributed data to the study only for ROIs in which they had electrode coverage; no minimum coverage criterion was imposed beyond inclusion in at least one ROI. The included ROIs were amygdala, anterior and posterior hippocampi, anterior cingulate cortex and orbitofrontal cortex. These regions were selected *a priori* based on their established roles in episodic memory formation, contextual event processing, and salience detection, processes thought to contribute to the encoding of event boundaries (Ding, Chen et al. 2006, Seth, Barrett et al. 2015). Across participants, each contributed data from three or more ROIs, depending on clinical implantation patterns. Regions used for statistical analysis required data from six or more contributing participants to be included to ensure sufficient statistical robustness.

Electrode localization was performed using YAEL (Wang, Magnotti et al. 2023), a pipeline that performs semi-automated electrode reconstruction from post-implantation CT and pre-implantation MRI based on Freesurfer imaging outputs. Individual T1-weighted MRI volumes were reconstructed using FreeSurfer (Fischl 2012) to generate subject-specific cortical surface models and subcortical parcellations. Post-implantation CT images were co-registered to the pre-implantation MRI for each subject. After electrode contact positions were confirmed, they were assigned to anatomical regions based on the co-registered Freesurfer segmentations. Contact assignments were visually verified by an experienced clinician for each subject.

Pseudocontacts were included in analyses if they were localized to one of the ten predefined bilateral ROIs. Contacts falling outside these ROIs, within white matter, or at the boundary between regions were excluded. Of the 89 total bipolar electrode pseudocontacts across all participants, 51% met these inclusion criteria and were retained for analysis. Participants contributed data only for ROIs in which they had at least one valid contact; no minimum contact number per ROI was imposed (see Table S1 for subject and pseudocontact contribution, and Table S2 for the number of included contacts per patient by anatomical region, and Figure S3 for electrode localization).

#### Clinical Seizure and Epileptiform Activity Review

For each patient, clinical records were reviewed to determine seizure activity and interictal epileptiform discharge rates in the 24 hours surrounding the experimental session. No patient experienced a seizure during the task itself; two patients had seizures earlier the same day, both concluding more than one hour before testing began. Interictal epileptiform discharges were observed in one or more analysis regions of interest in a subset of patients, with discharge rates ranging from less than 1 to 60 events per minute. To assess whether these findings influenced our central result, we repeated the group-level community detection analysis excluding the patient with the highest region-of-interest discharge burden, and separately excluding the two patients with same-day seizures; the cross-hemispheric, low-modularity community structure during Hits was preserved under both exclusions (see Discussion).

#### sEEG Signal Preprocessing

Raw sEEG signals were preprocessed prior to connectivity estimation. For each electrode included in the study based on the localization process, a notch filter was applied at 60 Hz to suppress line noise from electrical interference. Following filtering, signals were bipolar referenced by computing the difference between adjacent contacts along each electrode shaft for included contacts, effectively canceling common-mode noise and minimizing volume conduction artifacts while preserving local neural signals.

Bipolar referenced recordings data were segmented into trial-based epochs time-locked to stimulus onset during the Learn and Recognition phases of the task. Each epoch spanned from 200 ms prior to stimulus onset to 800 ms post-stimulus. A Hanning taper of 125 samples (250 ms at 500 Hz) was applied to each sliding window segment prior to autoregressive model fitting, reducing spectral leakage and edge effects (Kopal, Hlinka et al. 2023).

### Functional connectivity

#### Preprocessing

For each recording pseudocontact, the mean signal across trials was computed at each time point and subtracted from individual trials (ensemble mean subtraction). This procedure removes stimulus-locked, phase-consistent evoked responses while preserving trial-to-trial variability (induced activity). Removing these shared evoked components helps prevent spurious Granger causality estimates that can arise when multiple signals are driven by the same time-locked input rather than true inter-regional interactions (Ding, Chen et al. 2006). In addition, this step improves the approximate stationarity of the signals, which is an important assumption for autoregressive (AR) modeling, and was therefore applied alongside ensemble mean subtraction prior to connectivity estimation.

#### ROI Mapping and Group-Level Matrix Construction

Because electrode implantation varied across participants based on clinical considerations, individual channel-level connectivity estimates were mapped to a common anatomical framework to enable group-level analyses. Therefore, subjects contributed recording contacts to analyses only for ROIs in which they had electrode coverage; no minimum coverage criterion was imposed beyond inclusion in at least one ROI.

#### Sliding-window Granger Causality

For each subject, Granger causality (GC) values were averaged across all contact pairs, within a subject, belonging to the same ROI-to-ROI combination, yielding subject-level ROI connectivity matrices for each condition and time window. These subject-level matrices were then averaged across participants to generate condition-specific group-level matrices (Learn, Recog, and then further dichotomized into Hits and Correct Rejections). Directed connectivity was quantified using a bivariate Granger causality (GC) framework implemented in a sliding-window manner. GC estimates were computed using a window length of 250 ms, advanced in steps of 16 ms, yielding a time-resolved measure of directed inter-regional influence across each trial.

Within each window, bivariate autoregressive (AR) models of order M = 15 were fit to all channel pairs; previous work has shown model orders between 5 and 15 have shown no difference in results (Kopal, Hlinka et al. 2023). Model order specifies the number of past time lags (each separated by one sample) included in the autoregressive prediction, determining the temporal depth of the causal model. At a sampling rate of [SR] Hz, order 15 corresponds to approximately [15/SR x 1000] ms of history. Model order was selected to balance temporal resolution with model stability and has been shown to adequately capture short-lag dependencies in intracranial electrophysiological data (Kopal, Hlinka et al. 2023).

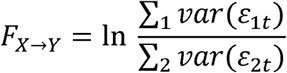

Directed GC from channel *X* -> *Y* was defined as the logarithmic ratio of the residual error variances between the reduced and full models:

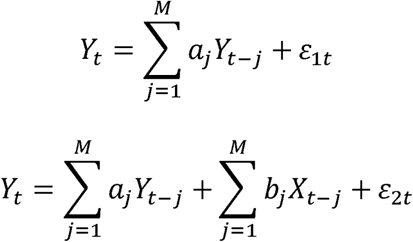

The GC F-statistic is defined as the logarithmic ratio of the reduced model to the full model. *X* and *Y* are recorded time series, *a* and *b* are parameters to the autoregressive model that correspond to the time lag *j*, and ɛ is the residual of the autonomous or coupled system. *t* is time and *M* is model order.

This measure reflects the extent to which past activity in channel *X* improves prediction of channel *Y* beyond what can be explained by *Y*’s own history. GC values were computed independently for each sliding window and trial, yielding time-resolved, direction-specific connectivity profiles (Granger 1969, Kopal, Hlinka et al. 2023). As GC values are defined to be greater than or equal to zero, any negative values that may be caused by computational instability were set to zero.

### Network construction and community detection

Group-level connectivity matrices were treated as weighted, directed networks, with ROIs as nodes and Granger causality (GC) values as edges. Community structure was identified using the Louvain modularity optimization algorithm (Blondel, Guillaume et al. 2008), applied independently to each sliding time window to capture the temporal evolution of network organization.

Because the Louvain algorithm is stochastic and can yield different solutions across runs (a property known as degeneracy), the algorithm was repeated 10,000 times per time window with random initializations. To obtain a stable estimate of community structure, the partition achieving the highest modularity Q across all 10,000 runs was selected as the representative solution for that window. This approach reduces sensitivity to random initialization and yields a reproducible representation of network organization.

Negative edge weights were handled asymmetrically following established procedures (Rubinov and Sporns 2011), allowing anti-correlated interactions to contribute to modular structure without dominating the optimization. The resulting time-resolved community assignments labeled each ROI according to its functional module at each time window. To maintain consistent community labels across time, assignments were aligned between adjacent windows using a greedy maximum-overlap algorithm: at each window, community labels were remapped to maximize the number of nodes sharing the same label as the preceding window’s partition. This automated procedure replaces the manual alignment used in Kopal et al. (Kopal, Hlinka et al. 2023) and produces equivalent results for stable two-community partitions. For downstream analyses, a single reference partition (M_final) was defined as the modal community assignment across all time windows, and GC strength was subsequently evaluated within these fixed community definitions over time.

Community labels (e.g., Community 1, Community 2) are assigned arbitrarily by the algorithm and therefore are not directly comparable across conditions. To enable consistent interpretation, communities were defined based on their anatomical composition rather than numeric label.

#### Dynamic Graph-Theoretic Metrics

To characterize the temporal evolution of network topology, two complementary graph-theoretic metrics were computed across sliding windows using the Brain Connectivity Toolbox (Rubinov and Sporns 2010). We computed network modularity (Q) to quantify the degree to which the network was organized into the communities identified by Louvain clustering, reflecting network segregation. For this metric, a higher Q indicates stronger within-community relative to between-community connectivity, meaning that brain regions within the same community interact more strongly with each other than with regions in other communities. In functional terms, this reflects a more segregated network organization, where communication is concentrated within specialized groups of regions rather than distributed across the whole network.

The second metric computed for each condition and community was the mean within-community GC strength, estimated as the average directed GC across all within-community ROI pairs at each time window. This provides a time-resolved measure of how strongly regions within a given community are interacting over time.

To quantify hemispheric lateralization of within-community connectivity, a hemisphere asymmetry index (AI) was computed for each community at each time window: AI = (R − L) / (R + L), where R and L denote the mean directed GC among right-hemisphere and left-hemisphere nodes within the community, respectively. Values range from −1 (fully left-dominant) to +1 (fully right-dominant). Significance was assessed using a sign-permutation bootstrap test against the null hypothesis of zero mean AI (10,000 permutations).

#### Hub Region Analysis

Hub regions were identified using three complementary centrality measures computed on time-averaged directed Granger causality (GC) matrices using MATLAB’s built-in graph analysis functions (The MathWorks 2025): degree centrality, betweenness centrality, and PageRank centrality (Brin and Page 1998). For degree centrality, nodes were thresholded at the top 10% of edge weights and combined in-degree and out-degree were computed. Betweenness centrality was defined as the fraction of shortest paths in the network that pass through a given node, capturing its role in mediating communication between regions. PageRank centrality was computed as an iterative measure that assigns higher importance to nodes receiving connections from other highly connected nodes.

Regions were classified as hubs if they consistently ranked within the top 20^th^ percentile of PageRank centrality within a given condition, with degree and betweenness reported as supplementary characterization. These metrics capture complementary aspects of hub-like influence: degree centrality reflects the number of connections a region has, betweenness centrality identifies regions that act as bridges between communities, and PageRank highlights regions that are influential based on both the number and importance of their connections. Hub region comparisons between Hits and Correct Rejections were used to characterize condition-specific differences in network organization.

### Statistical Analysis

#### Baseline Comparison

Statistical significance of community GC strength and global efficiency relative to the pre-stimulus baseline was assessed using a within-subject permutation test. For each condition, community, and post-stimulus timepoint *t*, the observed difference between each subject’s GC at *t* and their own pre-stimulus baseline mean was computed, defined as the average across all sliding windows occurring during the pre-stimulus interval spanning the full inter-trial interval (ITI), approximately –875 ms to –25 ms relative to stimulus onset, yielding approximately 47 baseline windows. This extended baseline replaces the four-window pre-stimulus estimate used in the original submission and provides a substantially more stable null reference.

A null distribution was generated by randomly shuffling, within each subject, the assignment of which observation served as the “post-stimulus” value versus baseline values, repeated across 10,000 permutations. The group-level test statistic (mean across subjects of the permuted difference) was compared against this null distribution, and a one-sided *p*-value was obtained. FDR correction was applied across post-stimulus timepoints only (Benjamini–Hochberg), and significance was determined at p_FDR < 0.05. Only timepoints ≥ 25 ms post-stimulus were included in reported significance maps to avoid contamination from the transition between pre- and post-stimulus windows.

This test required a minimum of six subjects with valid data for a given community. Community 2 during Hits, comprising left prefrontal and right MTL regions (LAC, LOF, RAH, RAMY, RPH), had bilateral coverage in only 2 of 7 subjects and was therefore excluded from subject-level significance testing; its dynamics are reported descriptively from the group meta-matrix.

#### Modularity Testing

The significance of observed modularity Q values relative to baseline was assessed using a multivariate frequency shuffling (MVFS) surrogate test (Kopal, Hlinka et al. 2023). For each surrogate iteration, GC timeseries were randomized independently for each channel pair and each subject, preserving the spectral structure of the signal while randomizing the temporal relationship. The surrogate GC matrices were used to rebuild the group-level meta-matrix on each permutation, and the Louvain algorithm was rerun to extract surrogate Q time courses. This procedure was repeated 1,000 times, yielding a null distribution of Q values at each timepoint that reflects the full variance introduced by network reconstruction, not merely phase randomization of the scalar Q timeseries. Observed Q at each post-baseline timepoint was compared against the surrogate distribution after subtracting the respective baseline means, and FDR correction was applied across timepoints. Timepoints ≥ 25 ms post-stimulus were included in reported significance maps.

To further validate that the observed modularity reflects genuine stimulus-related network organization rather than something spurious, we implemented a circular shift validation in which each channel pair’s GC timeseries was circularly shifted by a random lag, preserving autocorrelation structure and spectral content while destroying stimulus-locked temporal relationships. The full meta-matrix (variable described above) was rebuilt and Louvain clustering was rerun for each of 1,000 iterations per condition. At each post-stimulus timepoint, significance was assessed using the same one-sided percentile test and FDR correction procedure as the MVFS test. Results were concordant across both surrogate methods: no timepoints were significant under circular shift but not MVFS in any condition, confirming robustness of the reported modularity significance.

#### Hits vs. Correct Rejections Comparison

Condition differences in community GC strength between Hits and Correct Rejections were assessed at two levels. First, a trial-epoch-level permutation test compared per-subject mean GC strength (averaged across all post-stimulus timepoints) between conditions by pooling and randomly permuting the condition assignment (Hits vs. CR) within each subject across 10,000 permutations. A p-value was derived from the resulting null distribution, with significance accepted at α = 0.05 (one-tailed; observed value exceeding the 95th percentile of the permutation distribution). Second, a per-window permutation test was applied at each timepoint independently using the same label-permutation approach.

The temporal correlation between Hits and CR community strength sequences was assessed using Pearson correlation of the group-mean timeseries against a phase-randomized (MVFS) null model (1,000 surrogates), providing a test of whether the two conditions showed similar or divergent temporal dynamics independent of overall amplitude differences. Fewer surrogates were used here than for the Louvain consensus procedure (10,000) because MVFS surrogate distributions converge reliably at 1,000 iterations, whereas the stochastic nature of the Louvain algorithm requires a larger number of runs to achieve a stable consensus partition (Kopal, Hlinka et al. 2023).

#### Behavioral Correlations

Relationships between community GC strength and individual recognition memory performance (d’, a signal detection theory measure of discriminability computed as the standardized difference between hit rate and false alarm rate, with higher values indicating greater memory sensitivity) (i.e., a better ability to distinguish previously seen images from novel images independent of response bias) were assessed using Spearman correlations computed across subjects (n = 7). Two complementary analyses were conducted: (1) temporal correlations assessed, at each timepoint independently, the across-subject Spearman correlation between each subject’s mean community GC strength at that window and their session-level d’ score –yielding a time course of correlation coefficients reflecting whether subjects with stronger community connectivity at a given moment tended to show better overall recognition performance –with error correction applied across timepoints using a permutation approach to account for the high temporal autocorrelation between adjacent windows; and (2) summary correlations assessed the relationship between mean post-stimulus (0–300 ms) community strength and d’, with FDR correction across the four community *x* condition combinations tested. Given the small sample size, all behavioral correlation results are reported as exploratory and interpreted with appropriate caution. A permutation-based family-wise error correction controlling for the temporal autocorrelation structure of the timeseries was used for the temporal correlation analysis in place of standard FDR, as adjacent timepoints share approximately 94% of their data given the 250 ms window and 16 ms step.

#### Subject-Level Sensitivity Analysis

To evaluate the extent to which the group-level cross-hemispheric community finding was supported at the individual-patient level, we computed, for each patient independently, the mean Granger causality across all covered ipsilateral region pairs minus the mean GC across all covered cross-hemispheric region pairs, using only that patient’s own electrode contacts, separately for Hits and Correct Rejections and averaged over the post-stimulus window. Patients were included if they had at least one covered ipsilateral and one covered cross-hemispheric region pair.

## Results

### Behavioral Performance

Seven patients completed the recognition memory task with above-chance performance across all participants (Figure 2A, B). Group-level recognition sensitivity, the ability to distinguish previously seen from novel images, averaged d’ = 1.15 ± 0.27 (range: 0.77–1.43, Figure 2A). Mean overall accuracy was 69.6% ± 5.6%. Hit rates ranged from 0.30 to 0.78 (mean = 0.56) and false alarm rates from 0.06 to 0.32 (mean = 0.17). Response bias (criterion c) was computed as the negative average of the two z-transformed rates, with positive values indicating conservative response tendencies (bias toward ‘new’ responses) and negative values indicating liberal tendencies (biased toward ‘old’ responses; Figure 2C). Response bias was variable across subjects (mean c = 0.42 ± 0.39), with most participants adopting conservative response strategies, though bias and sensitivity were largely independent across individuals. These behavioral results confirm that all subjects performed the task successfully with meaningful trial counts in both Hit and Correct Rejection categories, providing a sufficient basis for condition-specific neural analyses.

**Figure 2.**
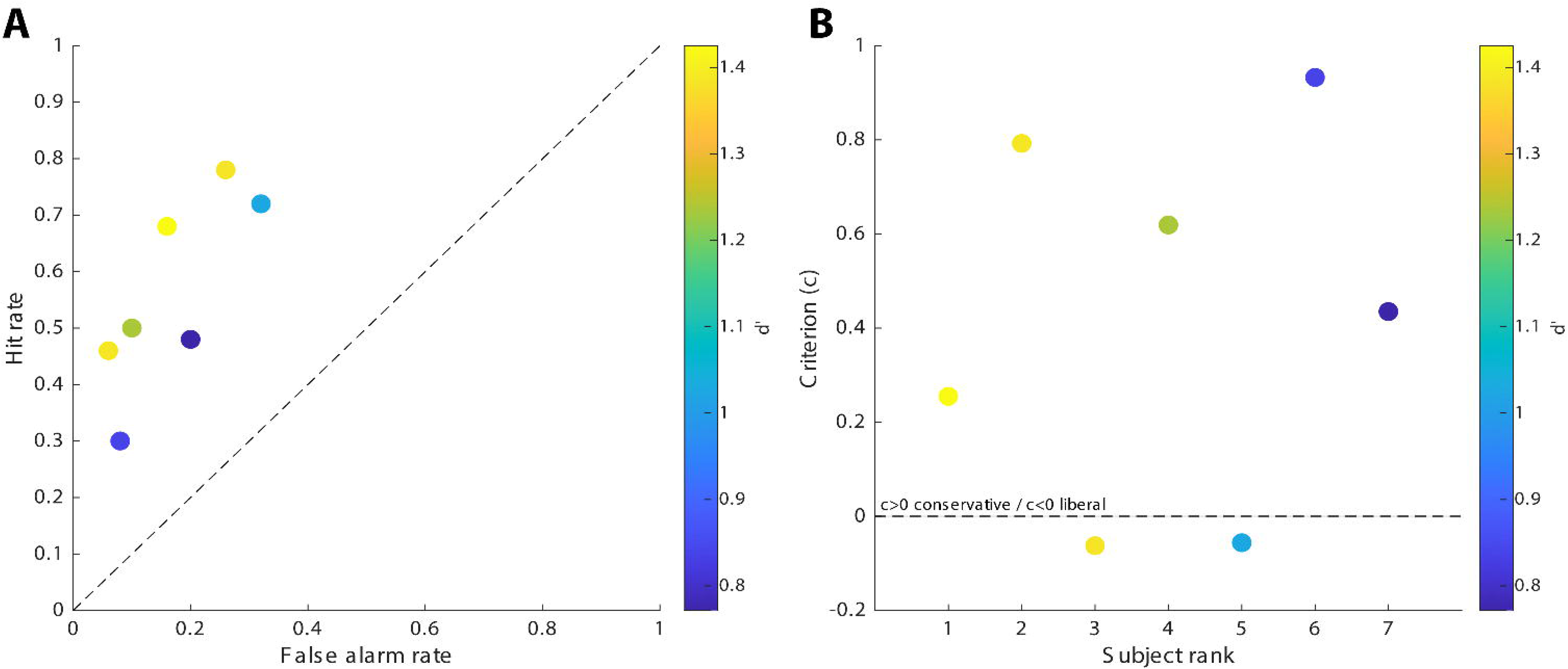
Behavioral performance across participants. (A) Recognition performance plotted in Receiver Operating Characteristic (ROC) space. Each point represents one participant, with false alarm rate on the x-axis and hit rate on the y-axis. Color encodes d’ (cool = lower, warm = higher sensitivity). The dashed diagonal represents chance performance (d’ = 0). All participants fall above the diagonal, confirming above-chance discrimination. (B) Response bias (criterion c) plotted by subject rank, colored by d-prime. Positive criterion values indicate conservative response strategies (tendency to respond “new”); negative values indicate liberal strategies (tendency to respond “old”). No systematic relationship between rank order and criterion was observed, suggesting that d’ variability reflects genuine differences in memory strength rather than response strategy.

Stimulus presentation duration was fixed at 1,000 ms across all trials. Mean reaction times were 926 ± 890 ms for Hits, 1,014 ± 1,143 ms for Correct Rejections, 1,216 ± 798 ms for False Alarms, and 960 ± 727 ms for Misses (mean ± SD across all trials and subjects). The large standard deviations reflect within-subject trial-level variability and between-subject differences in response speed. The post-stimulus analysis window (0–800 ms) captures neural dynamics that precede and likely contribute to the behavioral response rather than following it.

### Community Structure Differs Across Behavioral Conditions

To characterize large-scale network organization underlying directed connectivity, dynamic Louvain community detection was applied to condition-specific functional connectivity matrices derived from time-resolved GC estimates. Two dominant communities were identified consistently across all four conditions (Hits, Correct Rejections, Learn, Recognize), though community membership, modularity, and regional stability differed markedly as a function of behavioral outcome (Figures 3, 4).

**Figure 3.**
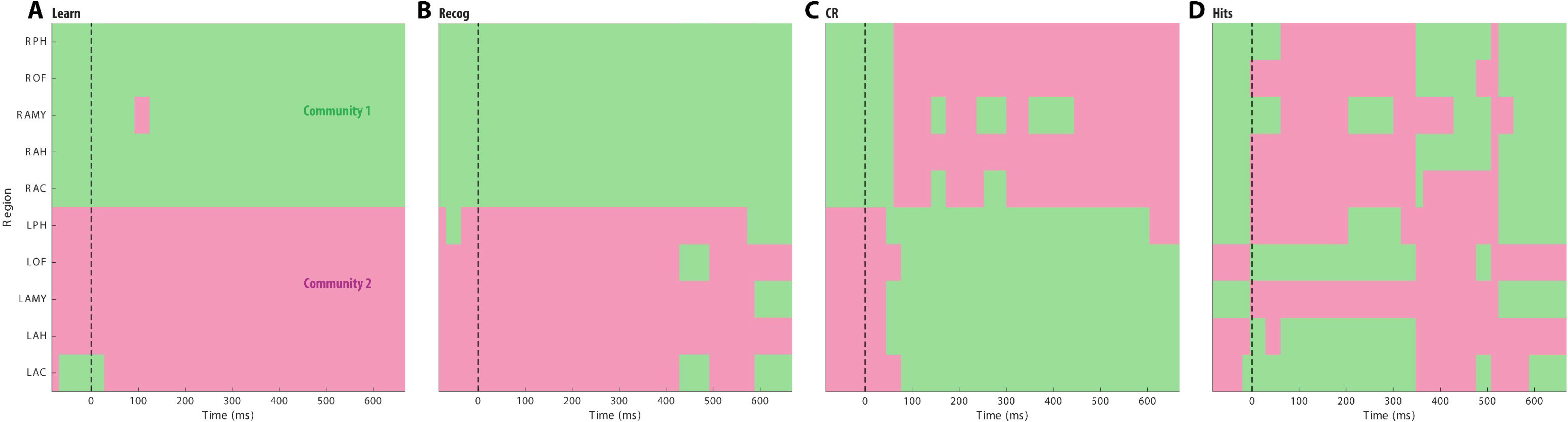
Dynamic community membership across the epoch. Each panel shows the time-resolved community assignment for each ROI across sliding windows for one condition, with Community 1 (green) and Community 2 (pink) color-coded. The dashed vertical line marks stimulus onset (0 ms). Hits are characterized by high regional instability, frequent switching between communities visible as fragmented color blocks, consistent with the low modularity observed in Figure 3. In contrast, baseline Learn and Recognition show near-complete stability after stimulus onset, with ROIs maintaining fixed community assignments across virtually the entire epoch. Correct Rejections show intermediate instability, with early transient switching in several left-hemisphere regions resolving to stable assignments by approximately 200 ms post-stimulus. The Hits panel uniquely shows sustained instability in bilateral MTL and prefrontal regions throughout the epoch, reflecting dynamic reorganization of large-scale network structure during successful recognition memory retrieval.

**Figure 4.**
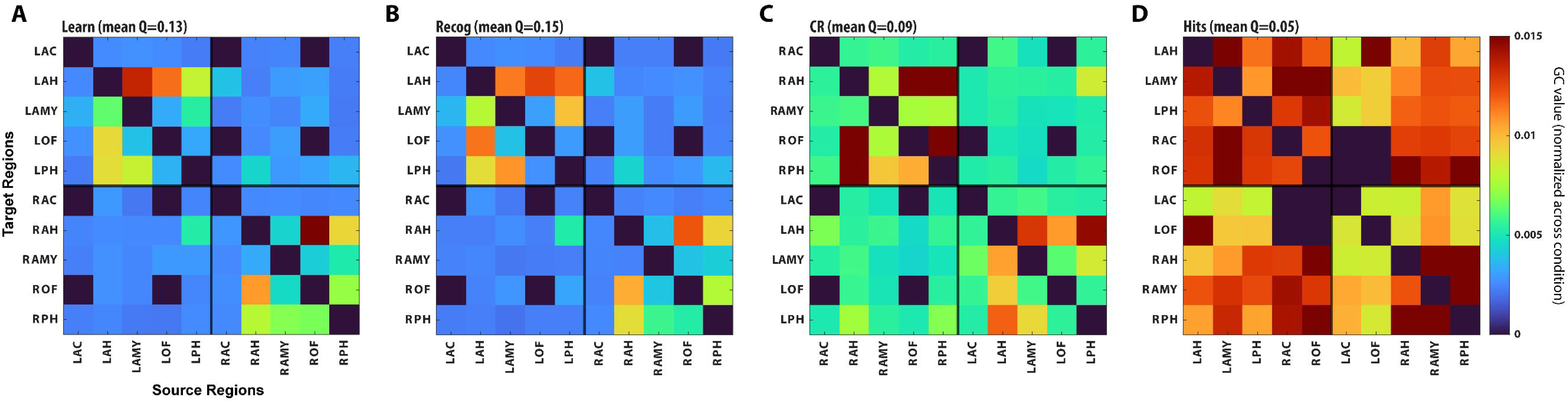
Time-averaged directed functional connectivity reordered by community structure. Heatmaps display mean Granger causality (GC) values averaged across the full epoch for each condition (Hits, Correct Rejections, Learn, Recognition), with ROIs reordered according to the Louvain community partition derived from that condition’s functional connectivity matrix. Black lines delineate community boundaries. Warm colors indicate stronger directed connectivity; cool colors indicate weaker connectivity. Mean modularity Q is reported above each panel. Hits (D) show the lowest modularity (Q = 0.05), reflecting diffuse connectivity that crosses community boundaries, whereas the complete Learn (A) and Recognition (B) periods maintain the highest modularity (Q = 0.13 and 0.15 respectively), consistent with an overall stable, segregated network organization. Notably, the Hits community partition groups left MTL regions (LAH, LAMY, LPH) with right prefrontal regions (RAC, ROF) into Community 1, a cross-hemispheric grouping absent in all other conditions. ROI abbreviations: L/R = left/right hemisphere; AC = anterior cingulate; AH = anterior hippocampus; AMY = amygdala; OF = orbitofrontal cortex; PH = posterior hippocampus.

During encoding (Learn) and recognition (Recog), community structure was highly stable and strongly hemispheric: one community comprised left-hemisphere ROIs (LAC, LOF, LAH, LPH, LAMY) and the other comprised right-hemisphere ROIs (RAC, ROF, RAH, RPH, RAMY), with regional instability indices near zero across regions. This bilateral segregation was consistent with the high modularity observed in these conditions (Q_Learn = 0.13, Q_Recog = 0.15) and reflects stable, lateralized network organization during non-retrieval memory portions. Community membership was essentially fixed after stimulus onset, with ROIs maintaining their assignments across virtually the entire epoch (Figure 3).

During Correct Rejections (CR), a similarly stable two-community structure emerged organized along hemispheric lines, with high modularity (Q_CR = 0.09) relative to Hits. Early transient switching in several left-hemisphere regions resolved to stable assignments by approximately 200 ms post-stimulus, after which community membership remained fixed for the remainder of the epoch. GC was greater in CR than Learn or Recog but not as high as in Hits.

In contrast, Hits were characterized by markedly reduced modularity (Q_Hits = 0.05, the lowest of all conditions), high regional instability, a cross-hemispheric community structure absent in all other conditions, and the highest overall GC. Community 1 during Hits comprised left MTL and right prefrontal regions (LAH, LPH, LAMY, RAC, ROF), while Community 2 comprised left prefrontal and right MTL regions (LAC, LOF, RAH, RPH, RAMY), a cross-hemispheric partition integrating medial temporal lobe with prefrontal cortex.

Sustained community instability was visible throughout the epoch in bilateral MTL and prefrontal regions (Figure 3), reflecting ongoing dynamic reorganization rather than a transient early response. This cross-hemispheric community structure, integrating left MTL with right prefrontal cortex and right MTL with left prefrontal cortex, was absent in all other conditions and constitutes a distinctive network signature of successful recognition memory retrieval.

### Hub Region Analysis and Hemispheric Lateralization

Hub regions were identified using degree, betweenness, and PageRank centrality applied to time-averaged directed GC matrices. Hub identity was dissociated between Hits and Correct Rejections, with no shared hub regions across conditions. During Hits, hubs were left-lateralized: left anterior hippocampus (LAH) in Community 1 and left orbitofrontal cortex (LOF) in Community 2. During Correct Rejections, hubs shifted to right-lateralized regions, with right amygdala (RAMY) emerging as the primary hub. During encoding (Learn), bilateral amygdala (LAH, RAH) served as hubs, and during general Recognition, left amygdala and left posterior hippocampus were prominent. The amygdala emerged as a hub region across all four conditions, underscoring its central role in the memory network regardless of behavioral outcome.

### Temporal Dynamics of Community-Level Connectivity

To assess how community-level directed connectivity evolved over time, mean within-community GC strength was computed across sliding windows for each condition and community, with significance assessed using a within-subject permutation test against each subject’s own pre-stimulus baseline (Figure 5).

**Figure 5.**
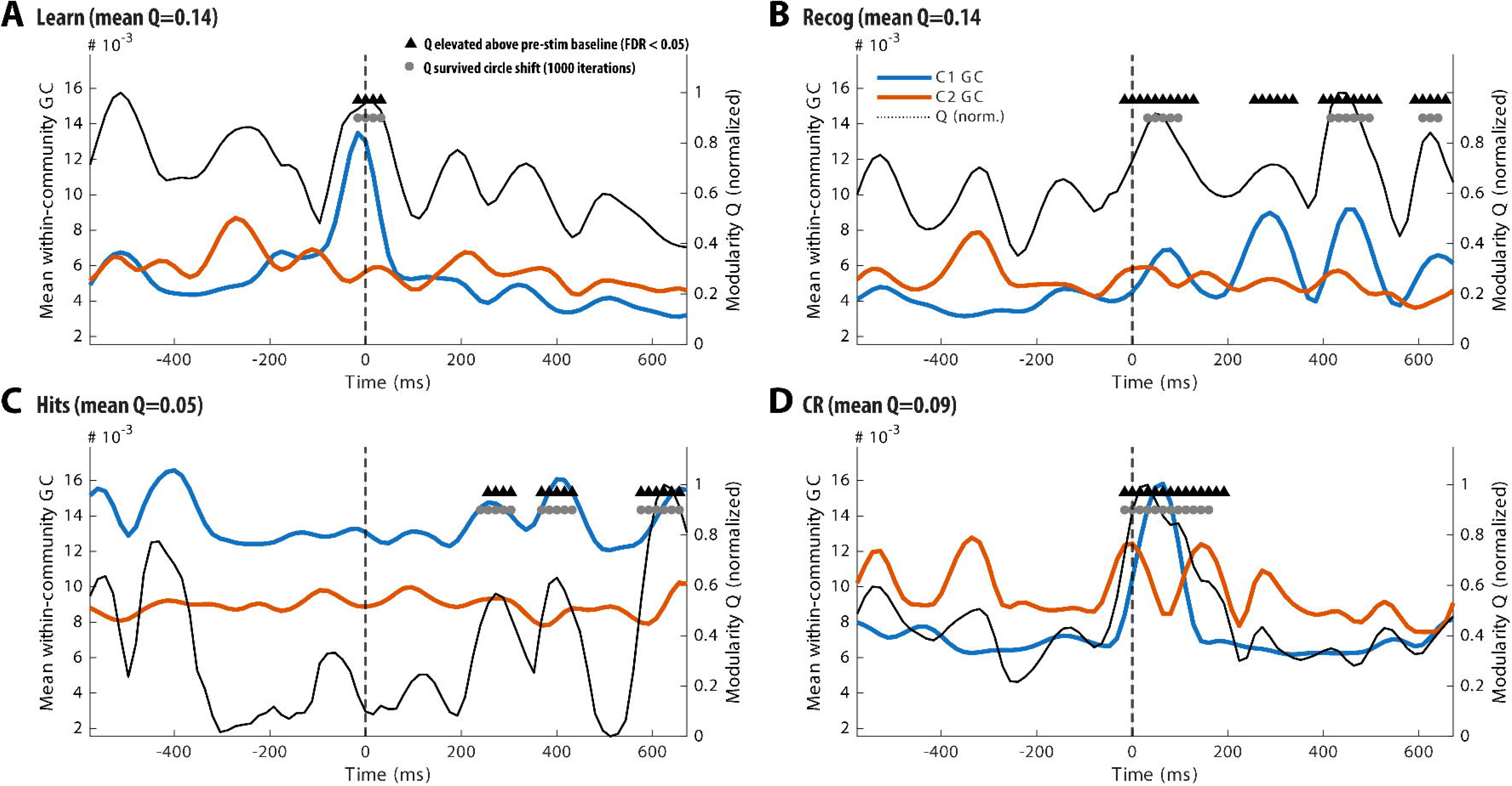
Community-level directed functional connectivity dynamics with anatomically consistent labeling and network modularity significance. Each panel shows mean within-community Granger causality (GC) strength over time aligned to stimulus onset (dashed vertical line, 0 ms), for two anatomically defined network communities: a left-hemisphere community (LAC, LAH, LAMY, LOF, LPH; solid blue) and a right-hemisphere community (RAC, ROF, RAH, RPH, RAMY; solid orange). However, in Hits, communities deviate from the pure hemispheric split –Community 1 (blue) comprises left MTL and right prefrontal regions (LAH, LPH, LAMY, RAC, ROF) and Community 2 (orange) comprises left prefrontal and right MTL regions (RAH, RPH, RAMY, LAC, LOF), reflecting the cross-hemispheric integration that distinguishes this condition. Community labels are mapped anatomically across conditions to enable direct cross-condition comparison. Note that the Louvain algorithm assigned arbitrary numeric labels independently per condition, which were remapped here by regional composition. The normalized modularity timecourse (Q, solid black, right axis), which reflects the degree of network segregation (i.e., stronger within-community relative to between-community connectivity), is overlaid for reference. Black triangles mark timepoints where Q was significantly elevated above pre-stimulus baseline (pFDR < 0.05, MVFS full-pipeline surrogate test, 1000 iterations). Gray circles indicate timepoints also surviving a circular shift validation (1,000 iterations). Condition-specific temporal dynamics and significance windows are described in the Results.

Community GC dynamics differed substantially across conditions in both temporal profile and significance (Figure 5). During CR, the right-hemisphere community (community 2) showed a prominent transient peak at stimulus onset followed by rapid decline, while the left-hemisphere community remained suppressed throughout the epoch; significant modularity was confined to the early post-stimulus window (32–192 ms), consistent with a brief segregated stimulus-driven response (Figure 5B). During Learn and Recog, GC dynamics were dominated by the left-hemisphere community with large oscillatory fluctuations throughout the epoch, and Recog showed the most sustained significant modularity of all conditions (32–672 ms; Figure 5C, D).

During Hits, both communities maintained elevated and broadly comparable GC strength throughout the epoch (Figure 5A). Neither community showed a significant baseline increase by the within-subject permutation test, which reflects genuinely high inter-subject variability in baseline connectivity rather than an absence of post-stimulus engagement: at the subject level, Community 1 showed a positive post-stimulus difference in 4 of 6 subjects, but one subject (MW16) with markedly elevated baseline GC (0.020, approximately 2.4x the group mean of 0.009) attenuated the group-level effect. Periods of significant modularity during Hits emerged late (256-304 ms, 368-432 ms) and clustered again near epoch end (576-672 ms), coinciding with a late rise in the right-hemisphere community GC.

The temporal correlation between Hits and CR community strength sequences was negative for both communities (left-hemisphere community: r = −0.17, p = 0.59; right-hemisphere community: r = −0.54, p = 0.13), confirming that the two conditions exhibit qualitatively divergent temporal dynamics rather than merely scaled versions of the same profile. Overall epoch-level GC strength was significantly higher during Hits than CR for the left-hemisphere community (permutation test, p < 0.0001), while the right-hemisphere community difference did not reach significance (p = 0.44), reflecting the more balanced inter-hemispheric profile during successful retrieval.

### Network Modularity and Integration Dynamics

Modularity dynamics differed across conditions in both magnitude and temporal profile. Modularity Q, which reflects the extent to which brain regions preferentially communicate within their own communities rather than across communities, provided a complementary characterization of network segregation dynamics, revealing condition-specific differences in both the timing and sustained significance of network organization. Hits maintained consistently low modularity across the entire post-stimulus epoch (Q_mean = 0.05), the lowest of all conditions, with significant periods emerging only after 256 ms. This delayed and sparse modularity during Hits is consistent with a persistently integrated network state following stimulus onset, in which community boundaries remain weak for several hundred milliseconds before partial segregation emerges late in the epoch.

CR showed significant modularity onset at 32 ms post stimulus, the earliest of all conditions, with a prominent early post-stimulus period of significant segregation (32–192 ms) coinciding with the transient right-hemisphere GC peak visible in Figure 5. This early modularity burst suggests that unsuccessful recognition is associated with an immediate segregated network response to the stimulus, in contrast to the sustained integration characterizing successful retrieval.

Learn and Recog conditions showed the highest sustained modularity (Q_mean = 0.13 and 0.15 respectively), consistent with their stable hemispheric community structure. Recog showed an extensive significant modularity across the entire epoch (onset 32 ms, sustained across multiple windows through 672 ms), reflecting the large oscillatory fluctuations in community GC visible in Figure 5.

False alarms had the lowest modularity (Q_mean = 0.04), and misses had a moderate modularity (Q_mean = 0.09) but the most extensive significant modularity across the epoch. Like Hits, false alarms had cross-hemispheric networks with the left and right amygdala serving as hubs (Figure S2). Misses followed the ipsilateral network pattern seen in Learn, overall Recog and CR (Figure S1).

To formally quantify differential suppression of cross-hemispheric relative to ipsilateral connectivity, linear mixed-effects models were fit separately for each condition, predicting z-scored Granger causality (GCz) from connection type (right → right, left → right, right → left, left → left categories), with a random intercept for subject. Ipsilateral connections (R → R, L → L) showed significantly higher GCz than cross-hemispheric connections (R → L, L → R) in all four conditions (all p = 0). However, the magnitude of this ipsilateral-cross-hemispheric gap differed systematically by condition, and was smallest during Hits and False Alarms and largest during Correct Rejections and Misses, mirroring the response-type dissociation (old vs. new judgment) described above rather than an accuracy-based dissociation. This trial-level analysis, incorporating all sliding-window GC estimates across subjects rather than the group meta-matrix used in the community-detection analyses, provides independent, convergent support for reduced hemispheric segregation during trials culminating in an “old” recognition judgment.

### Directed Network Topology at Key Post-Stimulus Timepoints

To characterize how directed network organization differed across conditions at specific moments, GC networks were reconstructed at two representative timepoints, an early window (∼115 ms) and a mid/late window (∼220 ms), based on prior work in these timepoints (Kopal, Hlinka et al. 2023), restricting analysis to the top 10% of edges by weight within each condition. Because thresholding was applied within-condition, edge presence reflects the dominant connectivity pattern for each condition rather than absolute inter-condition magnitude differences.

At the early timepoint (∼115 ms), Hits were uniquely characterized by prominent cross-hemispheric directed edges, most notably from right anterior cingulate (RAC) and right amygdala (RAMY) projecting to left amygdala (LAMY) and left anterior hippocampus (LAH). These cross-hemispheric connections were absent in all other conditions at this timepoint, which instead showed predominantly within-hemisphere ipsilateral connectivity concentrated in right-hemisphere chains (RAH -> RAMY -> RPH). This early cross-hemispheric connectivity during Hits indicates that integration of left MTL with right prefrontal networks begins within the first 100 ms of stimulus processing, substantially earlier than the community-level GC peak observed at approximately 420 ms.

At the mid/late timepoint (∼220 ms), all conditions converged on a predominantly right-hemisphere connectivity pattern, with the left hemisphere less engaged. Hits showed the sparsest network at this timepoint, consistent with the persistently low modularity during successful retrieval, while Correct Rejections maintained denser right-hemisphere coupling. The convergence across conditions at 220 ms, despite their divergence at 115 ms, suggests that the early cross-hemispheric integration uniquely observed during Hits is a transient phenomenon that dissipates as the network returns toward a more segregated configuration by the mid-epoch period.

### Subject-Level Sensitivity Analysis

To assess the individual-patient basis of the cross-hemispheric community finding, we repeated the analysis within each patient’s own electrode contacts independently, with no cross-subject pooling. For each patient with at least one covered ipsilateral and one covered cross-hemispheric region pair (6/7 patients; MW9 was excluded due to unilateral coverage), we computed the difference between mean ipsilateral and mean cross-hemispheric Granger causality during Hits and Correct Rejections. The two patients with right-prefrontal electrode coverage (MW12, MW16) both showed a smaller ipsilateral–cross-hemispheric gap during Hits than Correct Rejections, consistent with relatively greater cross-hemispheric engagement during successful recognition. The remaining four evaluable patients, whose right-hemisphere coverage was limited to posterior/medial structures (hippocampus, amygdala), did not show this pattern. This is consistent with an interpretation in which the effect is concentrated in and requires right-prefrontal coverage to detect at the individual level, and underscores that anatomical generalizability of the right-prefrontal component of this finding should be established in future work with broader coverage.

## Discussion

The present study characterized time-resolved directed functional connectivity during visual recognition memory using iEEG recordings from seven patients performing a new–old recognition task. By combining sliding-window Granger causality with dynamic community detection and graph-theoretic analysis, we identified a set of findings that collectively support a specific account of how large-scale network organization differs in successful recognition. Specifically, successful recognition (Hits) was associated with cross-hemispheric integration between left medial temporal and right prefrontal regions, reduced modularity, and elevated directed connectivity, a configuration absent in CR, Learn, and overall Recognize.

In the visual domain, recognition depends on the ventral visual stream, encompassing occipitotemporal regions involved in object identity and semantic representation (DiCarlo, Zoccolan et al. 2012, Grill-Spector and Weiner 2014). This ventral stream contribution is often lateralized, with right hemispheric regions showing disproportionate involvement in visual recognition tasks (Barr 1997, Guerin and Miller 2009). In addition, frontal and orbitofrontal cortices contribute to decision-making, confidence estimation, metacognitive monitoring, and response selection during judgments (Hoppstädter, Baeuchl et al. 2015, González, Zhang et al. 2024). Consistent with this framework, our findings reveal that large-scale topological organization—specifically, whether regions form cross-hemispheric communities—distinguishes Hits from other conditions, rather than regional activation alone.

This low modularity during Hits co-occurred with significantly elevated overall GC strength, indicating that while community boundaries were weak, total inter-regional communication was high. These data suggest successful retrieval reflects reduced segregation and elevated cross-regional signaling, enabling left MTL and right prefrontal regions to function as a unified network.

### Cross-Hemispheric Integration as a Network Signature of Successful Retrieval

The most striking finding was the emergence of a cross-hemispheric community during Hits that was relatively low in all other conditions. During successful recognition, left MTL regions (LAH, LAMY, LPH) were functionally integrated with right prefrontal regions (RAC, ROF) into a single network community, while left prefrontal and right MTL regions formed a second cross-hemispheric community. This bilateral, cross-hemispheric organization was replaced in CR, Learn, and Recog by a purely hemispheric partition—left hemisphere versus right hemisphere—with modularity values two to three times higher than during Hits and lower GC values.

This finding is consistent with theoretical accounts proposing that successful episodic and recognition memory retrieval depends on the coordinated engagement of medial temporal and prefrontal systems (Eichenbaum, Yonelinas et al. 2007, Rugg and Vilberg 2013). However, prior work has largely characterized this interaction in terms of overall connectivity strength or coherence (Staresina, Reber et al. 2019). Our results extend this literature by showing that during Hits, MTL and prefrontal regions are not merely correlated but functionally absorbed into a common community, sharing assignment across most post-stimulus windows.

The cross-hemispheric nature of this community was prominent. The left MTL has been associated with verbal and semantic memory encoding and retrieval (Golby, Poldrack et al. 2001), while right prefrontal cortex has been implicated in monitoring, verification, and post-retrieval evaluation processes (Hoppstädter, Baeuchl et al. 2015, González, Zhang et al. 2024). Their integration into a single community during Hits—but not Correct Rejections—suggests successful recognition requires both MTL activation and coupling with right prefrontal evaluative mechanisms (see Results, Subject-Level Sensitivity Analysis). Right-hemisphere nodes within this community showed stronger within-community connectivity than left-hemisphere nodes (mean AI = 0.33, p < 0.001), suggesting right prefrontal regions contribute more to intra-community drive than left MTL regions, consistent with top-down accounts of retrieval-mode engagement (Rugg and Curran 2007).

### Low Modularity as a Mechanism of Integration

The markedly reduced modularity during Hits (Q = 0.05 versus Q = 0.09–0.15 in other conditions) provides a mechanistic account of how cross-hemispheric integration is achieved. Low modularity reflects weak community boundaries, permitting cross-community signaling comparable to within-community signaling. During Hits, between-community GC approached or exceeded within-community GC from approximately 200 ms onward, a pattern absent in all other conditions.

This finding aligns with broader theoretical work linking cognitive flexibility and successful performance to transient reductions in network modularity (Bassett, Wymbs et al. 2011, Cohen and D’Esposito 2016). In the memory domain specifically, Westphal et al. (Westphal, Wang et al. 2017) demonstrated that successful episodic memory retrieval is associated with reduced global modularity and elevated frontoparietal-default mode coupling, predicting fewer false alarms. The present results extend this principle to the retrieval domain and to a sub-second timescale, demonstrating that the modularity reduction during Hits is likely not simply a tonic state difference but emerges dynamically, with significant modular periods not appearing until after 256 ms—well after the early sensory and familiarity-based responses that characterize the first 200 ms of recognition.

### Temporal Dissociation Between Hits and Correct Rejections

The temporal dynamics of community GC and modularity revealed a clear dissociation between Hits and Correct Rejections that extends beyond the epoch-level community structure differences. Correct Rejections were characterized by an early modularity burst coinciding with a transient right-hemisphere GC peak at stimulus onset, followed by rapid decay. This pattern reflects a brief, segregated stimulus-driven response in which right MTL–hippocampal networks reject novel stimuli without broader integration.

Hits, by contrast, showed no early modularity peak and maintained a persistently integrated network state from stimulus onset, with late-emerging modular periods only after 256 ms. This sustained integration likely reflects the time required to retrieve and evaluate the memory trace before partial re-segregation later in the epoch. The negative temporal correlation between Hits and CR community strength timecourses (Community 2: r = −0.54) further confirms that these conditions follow qualitatively divergent, not merely scaled, temporal trajectories.

The directed network topology analysis at specific timepoints converged with this interpretation. At ∼115 ms, Hits uniquely exhibited cross-hemispheric directed edges from right prefrontal to left MTL regions, suggesting that the integration process begins early—within the first 100 ms of stimulus processing—even though its community-level signature does not peak until ∼420 ms. This early cross-hemispheric connectivity, absent in other conditions, may reflect an initial top-down prefrontal signal relayed to medial temporal regions to initiate retrieval.

### Right-Hemisphere Dominance in Non-Retrieval Conditions

An unexpected finding was the consistency of right-hemisphere dominance across Correct Rejections, Learn, and Recog. In all three conditions, within-condition GC dynamics were dominated by the right-hemisphere community, with right amygdala and right posterior hippocampus serving as primary hub regions during CR. This is consistent with prior reports of right-hemisphere specialization for visual recognition in both neuropsychological (Barr 1997, Guerin and Miller 2009) and electrophysiological literatures (Barbeau, Taylor et al. 2008). However, that this dominance characterizes Correct Rejections rather than Hits suggests: the right MTL network may be more engaged during rejection of novel stimuli, while successful retrieval requires left MTL recruitment and cross-hemispheric integration.

### Limitations

Several limitations warrant consideration. First, the sample size of seven patients limits statistical power for subject-level analyses and precludes reliable brain-behavior correlations at conventional thresholds. In particular, only two of seven patients (MW12, MW16; Table S1) had electrodes in right frontal ROIs (RAC, ROF), meaning the right-prefrontal component of the cross-hemispheric community structure reported here is derived from a small subset of the cohort (see Results, Subject-Level Sensitivity Analysis). Second, electrode coverage was heterogeneous across patients, meaning the community-level dynamics for networks were estimated primarily from the group meta-matrix rather than consistent subject-level data. Importantly, because electrode location was clinically determined in this epilepsy population, structural brain networks may differ from non-epileptic populations; to minimize this confound, primary analyses focused on within-patient differences across memory states rather than connectivity between specific regions. Relatedly, interictal epileptiform activity was present in analysis regions in a subset of patients; a sensitivity analysis excluding the patient with the highest discharge burden and, separately, the two patients with same-day (pre-task) seizures preserved both the cross-hemispheric Hits community structure and the direction of the Q_Hits < Q_CR modularity difference in every scenario tested, though we cannot rule out subtler effects on connectivity estimates at the single-contact level. Third, bivariate GC cannot rule out shared input from a third, unmodeled region driving apparent pairwise interactions; however, bivariate approaches remain standard in the iEEG connectivity literature and have been validated against multivariate methods in comparable datasets (Kopal, Hlinka et al. 2023). Together, these findings point to a specific circuit-level mechanism in which right prefrontal signals integrate with left MTL memory representations during the first several hundred milliseconds of stimulus processing, distinguishing successful retrieval by network topology rather than connectivity amplitude. Future work with larger cohorts and broader coverage is needed to assess generalizability and determine whether this network reflects known memory systems or a novel sub-second organizational principle. Similarly, response-locked analyses, once larger per-subject trial counts permit reliable estimation at the individual level, could clarify the temporal relationship between this network reorganization and the behavioral response itself.

## Supporting information

Supplementary Analysis

## Declaration of Generative AI and AI-Assisted Technologies in the Manuscript Preparation Process

AI-assisted writing tools (Grammarly; ChatGPT GPT4/5, OpenAI) were used to assist with manuscript editing and revision. All scientific content, analyses, interpretations, and conclusions were generated and verified by the authors. AI assistance was limited to language editing and structural organization of the text.

## Acknowledgments

This work was supported by NIH grant U01NS117839 to J.T. and U.R.

---

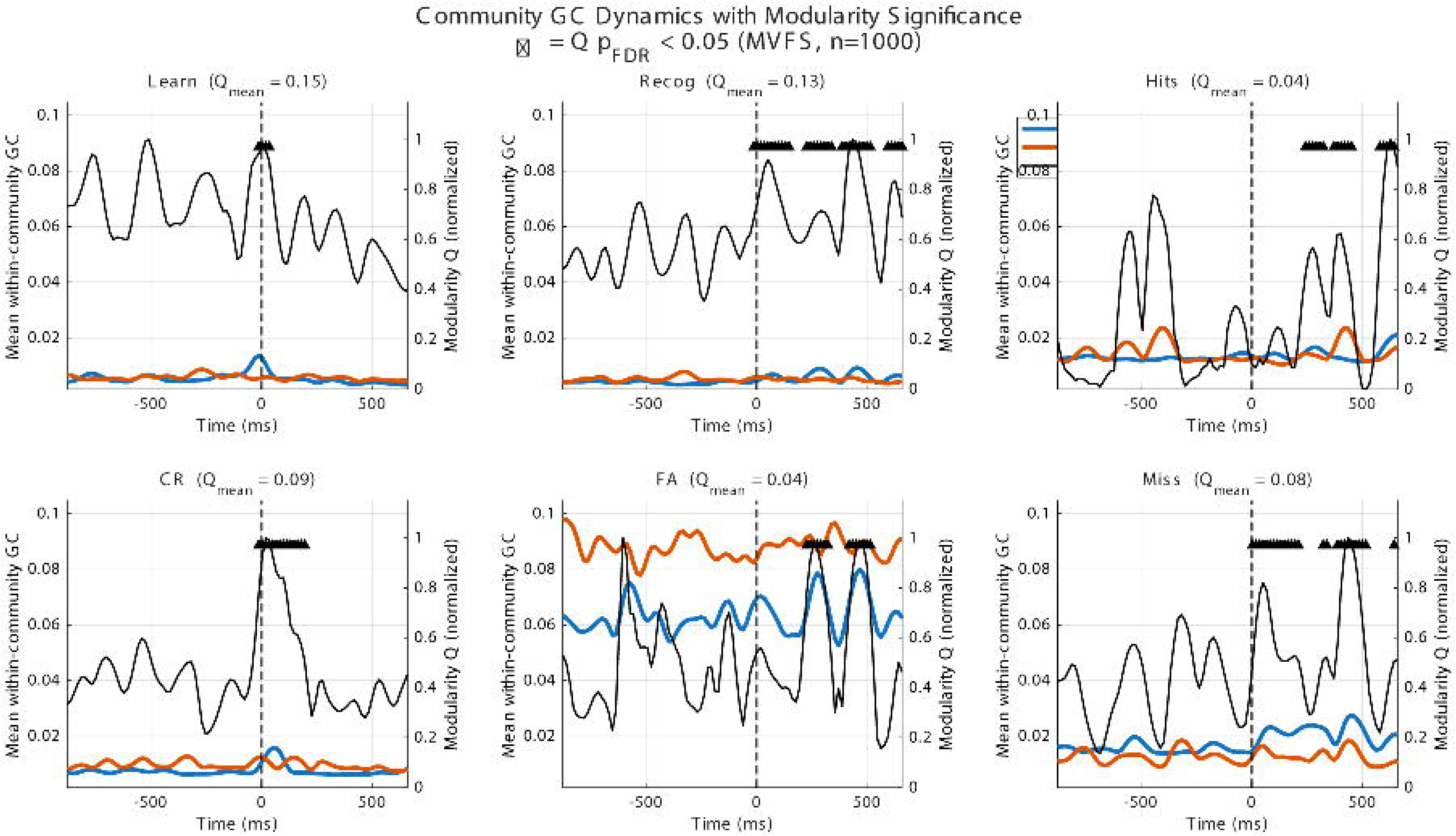

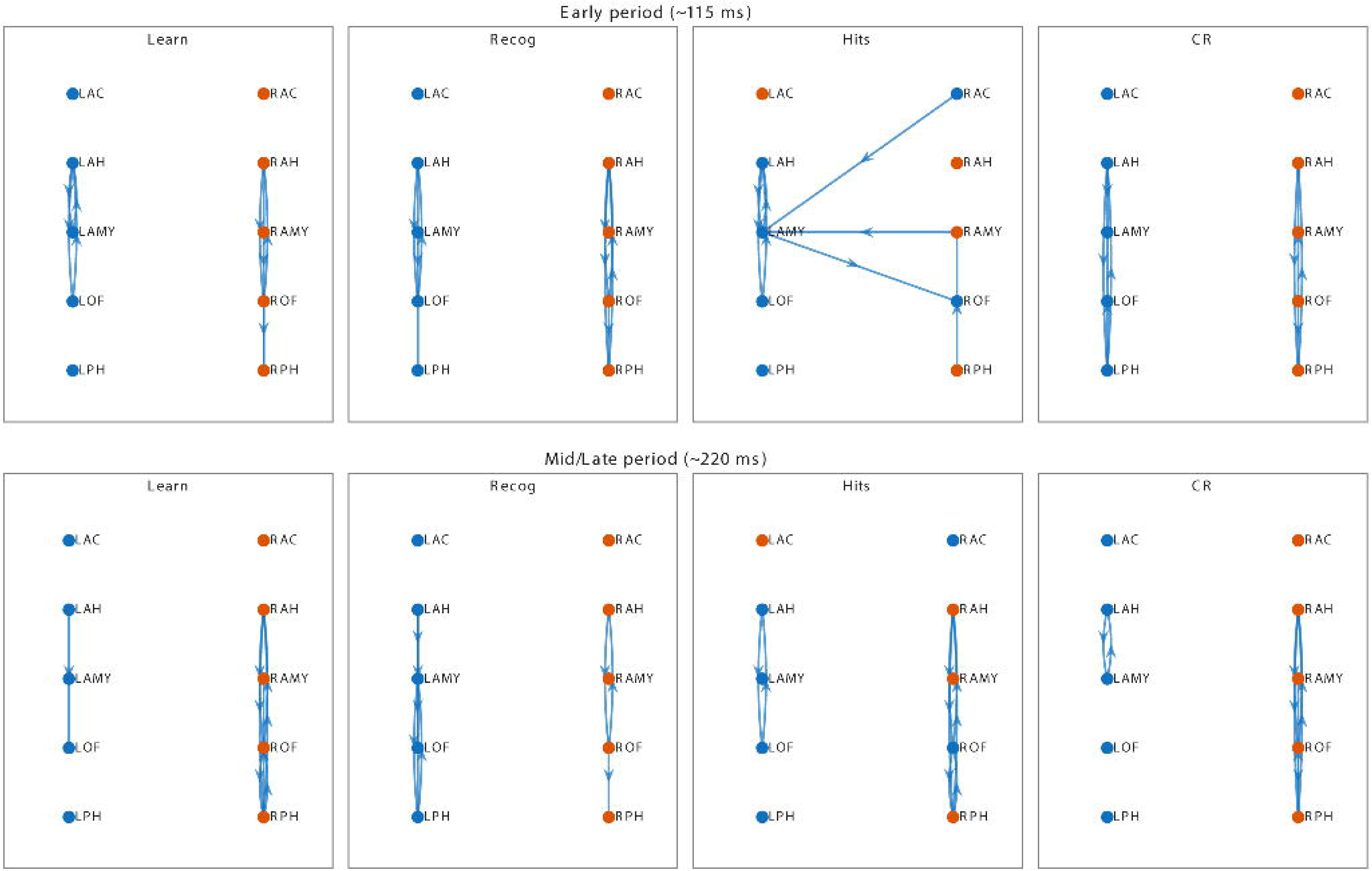

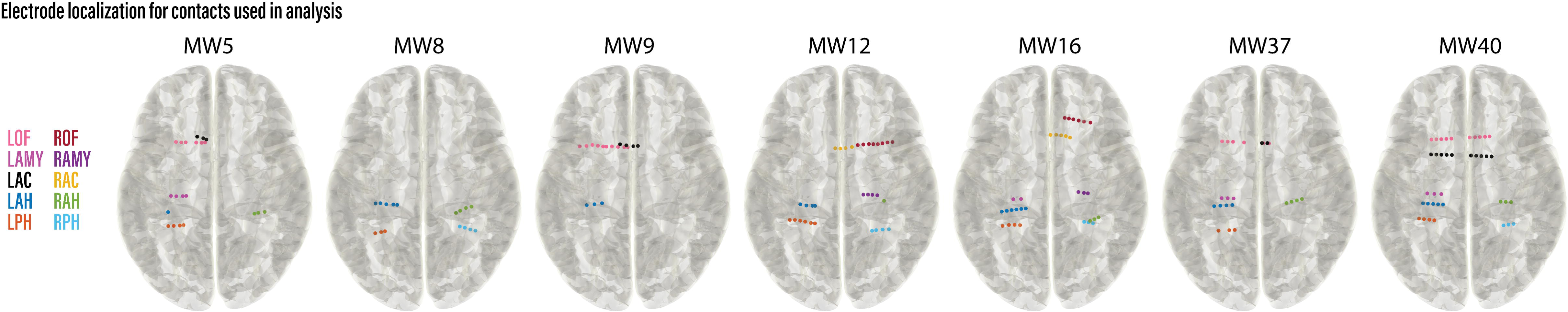

