## Supplementary Analysis for "Stable Network Motifs with Divergent Engagement Across Memory Outcomes"

| **Sex** | **Age** | **Dominant Hand** | **# Electrodes** | **% Contacts w/in ROI** |
| --- | --- | --- | --- | --- |
| F | 49 | R | 14 | 52 |
| F | 41 | L | 11 | 38 |
| F | 33 | R | 11 | 29 |
| M | 46 | R | 14 | 54 |
| M | 23 | R | 13 | 67 |
| F | 44 | R | 12 | 56 |
| M | 38 | R | 14 | 54 |

**Table S1.** Table summarizing pertinent characteristics from each patient. All patients have a medical history of epilepsy and are refractory to medication. During Phase II SEEG implantation, they performed a new-old delay task (depicted below). Because the electrode implantation varied among these patients, the final column shows what percent of all contacts on the electrodes were within our defined regions of interest, namely, bilateral orbitofrontal cortex, hippocampus, cingulate and amygdala. Out of all patients, ~51% of all contacts were contained in these regions of interest.

| **Subject** | **Anterior Cingulate** | **Amygdala** | **Anterior Hippocampus** | **Posterior Hippocampus** | **Orbitofrontal** |
| --- | --- | --- | --- | --- | --- |
| 1 | 3 (25%) | 4 (20%) | 4 (17%) | 4 (33%) | 6 (43%) |
| 2 | - | - | 9 (30%) | 7 (25%) | - |
| 3 | 4 (25%) | - | 3 (38%) | - | 10 (63%) |
| 4 | 4 (25%) | 4 (29%) | 5 (18%) | 10 (33%) | 9 (56%) |
| 5 | 5 (28%) | 5 (22%) | 9 (33%) | 7 (23%) | 6 (33%) |
| 6 | 3 (17%) | 3 (33%) | 8 (38%) | 3 (20%) | 6 (40%) |
| 7 | 10 (67%) | 3 (33%) | 8 (33%) | 7 (29%) | 10 (67%) |

**Table S2.** Included contacts per patient by anatomical region of interest. For each patient, the number of contacts confirmed via anatomical reconstruction to lie within gray matter of each bilateral region of interest is shown. Percentages reflect the number of confirmed in-target contacts as a proportion of all contacts on the corresponding electrode lead(s) that were nominally labeled to that region prior to individual contact-level verification; not all contacts on a labeled lead were ultimately confirmed to lie within target gray matter (some fell in white matter, at regional boundaries, or in adjacent structures and were excluded from the numerator).

### False Alarms and Misses

To further characterize the specificity of the network signatures identified in the primary analysis, directed GC networks and community structure were also examined for false alarm (FA) trials (novel images incorrectly judged as old) and miss trials (previously seen images incorrectly judged as new). These error trial types provide a dissociation between the effects of behavioral outcome (correct vs. incorrect) and response type (old vs. new judgment).

Community structure analysis revealed that FA trials, like Hits, were characterized by cross-hemispheric community organization (Q_mean = 0.04), with Community 1 comprising a mixture of left and right hemisphere regions (LAMY, RAC, ROF, RAH, RPH) and Community 2 comprising the remaining cross-hemispheric set (LAC, LOF, LAH, LPH, RAMY). This cross-hemispheric integration during FA closely mirrored the Hits community structure, and FA exhibited the lowest modularity of all six conditions (Q = 0.04 vs. Q = 0.05 for Hits). Miss trials, by contrast, showed purely ipsilateral community organization identical in structure to Correct Rejections: Community 1 comprised left-hemisphere ROIs (LAC, LOF, LAH, LPH, LAMY) and Community 2 comprised right-hemisphere ROIs (RAC, ROF, RAH, RPH, RAMY), with Q_mean = 0.09.

This pattern reveals a dissociation: cross-hemispheric community structure was observed in both conditions involving an "old" recognition response, whether correct (Hits) or incorrect (FA), while ipsilateral hemispheric segregation characterized both conditions involving a "new" response, whether correct (CR) or incorrect (Miss). Community structure therefore tracked response type rather than response accuracy, suggesting that the bilateral network integration observed during Hits reflects a general neural signature of the recognition decision itself rather than a marker of successful memory retrieval per se.

This interpretation is further supported by the network snapshots at key post-stimulus timepoints (Figure S2). At the early timepoint (~115 ms), FA showed the most prominent cross-hemispheric connectivity pattern in the dataset, with left amygdala (LAMY) receiving convergent directed input from multiple right-hemisphere regions (RAC, ROF, RAH, RAMY), a more widespread cross-hemispheric hub pattern than observed during Hits at the same timepoint. By the mid/late timepoint (~220 ms), FA maintained cross-hemispheric edges while Hits connectivity became sparse, suggesting that the cross-hemispheric integration in FA may be more sustained or less efficiently resolved than in correct recognition. Miss trials showed a connectivity pattern closely resembling Correct Rejections at both timepoints, with predominantly right-hemisphere ipsilateral chains and minimal cross-hemispheric edges.

The modularity dynamics figure (Figure S1) further illustrates this dissociation: FA showed broadly significant modularity across the epoch despite its low mean Q, while Miss showed a pattern of significant modularity windows more similar to CR and Learn. Taken together, these findings suggest that the low-modularity cross-hemispheric network state is a correlate of the recognition memory decision process – the subjective sense of familiarity driving an "old" response – rather than a reliable marker of veridical memory retrieval. Whether this reflects genuine familiarity-based processing or a failure of recollection-based rejection in FA trials remains an open question for future work with larger trial numbers per subject.

**Figure S1. Network modularity dynamics for false alarm and miss trial types.** Each panel shows mean within-community Granger causality (GC) strength over time aligned to stimulus onset (dashed vertical line, 0 ms), for the left-hemisphere community (solid blue) and right-hemisphere community (solid orange), with the normalized modularity timecourse (Q, solid black, right axis) overlaid for reference. Black triangles mark timepoints where Q was significantly elevated above pre-stimulus baseline (p < 0.05 FDR-corrected, MVFS surrogate test). FA trials show a cross-hemispheric community structure with broadly significant modularity across the epoch despite low mean Q, consistent with the pattern observed in Hits. Miss trials exhibit a right-hemisphere-dominant GC profile with significant modularity windows more similar to Correct Rejections and Learn. ROI abbreviations as in Figure 3.

**Figure S2. Early vs mid/late networks, including FA and Miss.** As in Hits, FA shows a cross-hemispheric network in both early and mid/late-period networks. Only connections in the top 10% of GC strength were visualized. The color of the node indicates community assignment per Louvain clustering.

**Figure S3. Electrode localization for contacts used in analysis.** For all patients included in the study, this figure visualizes the location of each included contact used in analysis, and that was part of the described regions of interest.
